# Comparative profiling of sperm small RNAs in cnidarians highlights potential for paternal inheritance

**DOI:** 10.64898/2026.11.02.763747

**Authors:** Jill Ashey, Benjamin H. Glass, Nicholas S. Galambos, Colin C. Conine, Katie L. Barott

## Abstract

Small RNAs in mature sperm are crucial regulators of gene expression following fertilization during early development in bilaterians. However, the presence and functions of sperm-derived small RNAs remain poorly understood in early-diverging (i.e., non-bilaterian) metazoans. In this study, we characterize three classes of small RNAs—microRNAs (miRNAs), piwi-interacting RNAs (piRNAs), and tRNA-derived RNAs (tDRs)—in sperm from three phylogenetically and ecologically distinct species from the early-diverging phylum Cnidaria: the sea anemone *Nematostella vectensis*, the temperate stony coral *Astrangia poculata*, and the reef-building tropical coral *Acropora hyacinthus*. All three classes of small RNAs examined were present in sperm from these cnidarians. Most miRNAs were species-specific; however, target prediction analysis revealed enriched putative targets involved in conserved developmental and signaling pathways. piRNAs overlapped with transposable elements and genic regions, indicating a potential dual role in genomic maintenance and gene expression regulation. Characterization of tDRs demonstrated that 5’-terminal fragments constituted the primary structural type across the three species, driven by high accumulation of specific isoacceptors including 5’-tDR-GlyGCC. Together, these findings provide the first evidence that cnidarian sperm carry a rich repertoire of small RNAs, including both conserved and species-specific types. Their presence suggests that paternal small RNAs contribute important regulatory information to the zygote in cnidarians, implicating paternal small RNAs as possible mediators of epigenetic inheritance in this early-branching metazoan phylum.

## Introduction

The transmission of biological information across generations is the foundation of inheritance, adaptation, and speciation. While classical genetic theory posits that inherited traits are exclusively transmitted though nuclear DNA sequence variation, an expanding body of evidence demonstrates that organisms transmit non-genetic (i.e., epigenetic) molecular signals that shape offspring development, phenotype and fitness (Fitz-James and Cavalli 2022). These signals can be modulated by the external environment, making them important vectors of environmental information across generations (Ho and Burggren 2010; Norouzitallab et al. 2019). Among the molecular features capable of mediating epigenetic inheritance, small non-coding RNAs (small ncRNAs) have emerged as important drivers of intergenerational information transfer (Sharma 2019; Fitz-James and Cavalli 2022).

Small ncRNAs encompass several diverse classes, including microRNAs (miRNAs), piwi-interacting RNAs (piRNAs), and tRNA-derived RNAs (tDRs), which exhibit distinct accumulation patterns across cell types, particularly in the germline. Canonical miRNAs (∼20–25 nt) associate with Argonaute proteins to regulate post-transcriptional gene expression via mRNA degradation or translational repression (Kaikkonen et al. 2011). In contrast, piRNAs (∼24–32 nt) regulate genomic stability by associating with Piwi proteins that silence transposable elements through transcriptional and post-transcriptional mechanisms (Czech et al. 2018).

However, recent evidence demonstrates that piRNAs can also regulate gene expression, albeit via distinct mechanisms from miRNAs (Rojas-Ríos and Simonelig 2018). tDRs (∼30–40 nt) are an emerging class of small ncRNAs that are generated by enzymatic cleavage of mature tRNAs (Akiyama and Ivanov 2023). Depending on the cleavage site relative to the mature tRNA structure, tDRs can be classified into distinct structural types (5’, internal, or 3’), which interact with RNA-binding proteins and translational machinery to regulate gene expression (Su et al. 2020; Akiyama and Ivanov 2023; Muthukumar et al. 2024). Together, these small ncRNA pathways form sophisticated gene regulatory networks that are active not only within somatic tissues, but also within mature germ cells.

Historically, the paternal contribution to epigenetic inheritance was assumed to be negligible owing to the small size and simple structure of sperm cells (Krawetz 2005). Mature sperm were long viewed as transcriptionally silent carriers, used solely as a means to deliver the paternal haploid genome to the oocyte. However, recent work has dismantled this paradigm. Mature sperm are now known to harbor a complex payload of small regulatory RNAs packaged during spermatogenesis (Sharma et al. 2016; Conine et al. 2018). Rather than residual transcriptional debris, sperm small RNAs constitute active regulatory molecules that can be shaped by paternal environmental exposures including thermal stress (Trigg et al. 2024), nutritional shifts (Chen et al. 2016; Tomar et al. 2024), and other physiological disruptions (Crews et al. 2012; Rodgers et al. 2013, 2015; Galambos et al. 2026).

Despite advances in our understanding of sperm-mediated epigenetic inheritance via small RNAs (Lee and Conine 2022; Naveed et al. 2025), current knowledge is based almost exclusively in bilaterian model organisms. This narrow taxonomic focus leaves a fundamental gap in understanding the evolutionary biology of epigenetic inheritance: is sperm-borne small RNA inheritance a recent bilaterian innovation or does it represent an ancient metazoan strategy conserved across early-diverging lineages? Evaluating the possible evolutionary origins of paternal small RNA inheritance requires profiling sperm small RNAs in non-bilaterian phyla that represent the base of the animal tree.

The phylum Cnidaria (e.g., sea anemones and corals) is the sister group to Bilateria (Miller et al. 2005), and thus provides an ideal evolutionary model for addressing this gap. Beyond their evolutionary relevance, cnidarians, particularly reef-building scleractinian corals, are ecologically and economically important species that are increasingly threatened by anthropogenic climate change (Hughes et al. 2017). Because many reef-building coral species are long-lived and reproduce only a few times annually, their capacity to adapt to climate change through natural selection acting on standing genetic variation is constrained. Non-genetic mechanisms, such as sperm-mediated small RNA inheritance, offer a conduit for rapid intergenerational plasticity, potentially allowing parental environmental experiences to shape offspring phenotypes and facilitate cross-generational acclimatization (Torda et al. 2017).

Therefore, investigating sperm small RNA profiles from cnidarians could illuminate the evolutionary origins of paternal epigenetic inheritance and its potential role in mediating cross-generational acclimatization to rapid environmental change in threatened species (Obura et al. 2026).

Here, we present the first comparative profiling of the sperm small RNA repertoire (including miRNAs, piRNAs, and tDRs) across three phylogenetically and ecologically distinct cnidarians: the sea anemone *Nematostella vectensis*, the temperate stony coral *Astrangia poculata,* and the reef-building tropical coral *Acropora hyacinthus*. Spanning over 600 million years of cnidarian evolution (Kayal et al. 2018), these species encompass diverse life history strategies, symbiotic modes, and ecological niches. By characterizing their sperm miRNAs, piRNAs, and tDRs, we establish the foundational small RNA landscape of cnidarian sperm. Further, we identify conserved processes likely targeted upon fertilization, and provide a foundation for future functional and evolutionary studies regarding epigenetic inheritance in these important and imperiled taxa.

## Materials and Methods

### Spawning and sperm sample collection

For *Astrangia poculata* (gonochoric broadcast spawners; Ashey et al. 2025; Figure 1A), spawning was induced and sperm samples were collected as described in Glass et al. (2023). Briefly, adult colonies (∼5–7 cm in diameter) were collected with a hammer and chisel at a depth of ∼3 m from Narragansett Bay, RI, USA (41.49231, −71.41883) in August 2022. Colonies were transported in coolers containing seawater to the University of Rhode Island, and individually placed in bowls of seawater (∼50 mL) sourced from Narrangansett Bay. Spawning was induced by exposing corals to an acute heat treatment (ramp from 22°C to 31°C over 30 min; Borbee et al. 2026). Temperature was controlled using an aquarium heater (Aqueon Pro Heater, AP300W-H) placed in a heated water bath that surrounded the bowls containing individual corals. Corals were returned to ambient temperature (22°C) following the acute exposure, and spawned within 1–8 h of heat exposure. Sperm were collected from individual male colonies immediately upon spawning. For collection, seawater containing sperm was first passed through a 100 µm cell stainer, then concentrated sperm pellets were created by repeatedly centrifuging (2–3 spins) 1 mL aliquots of the resulting filtrate (∼1 mL) at 1500 x *g* for 5 min, with the supernatant removed and replaced with filtrate in the same tube between each spin. After the final spin, seawater was removed, and pellets (∼50–100 µL) were stored at –80°C until further processing.

**Figure 1.**
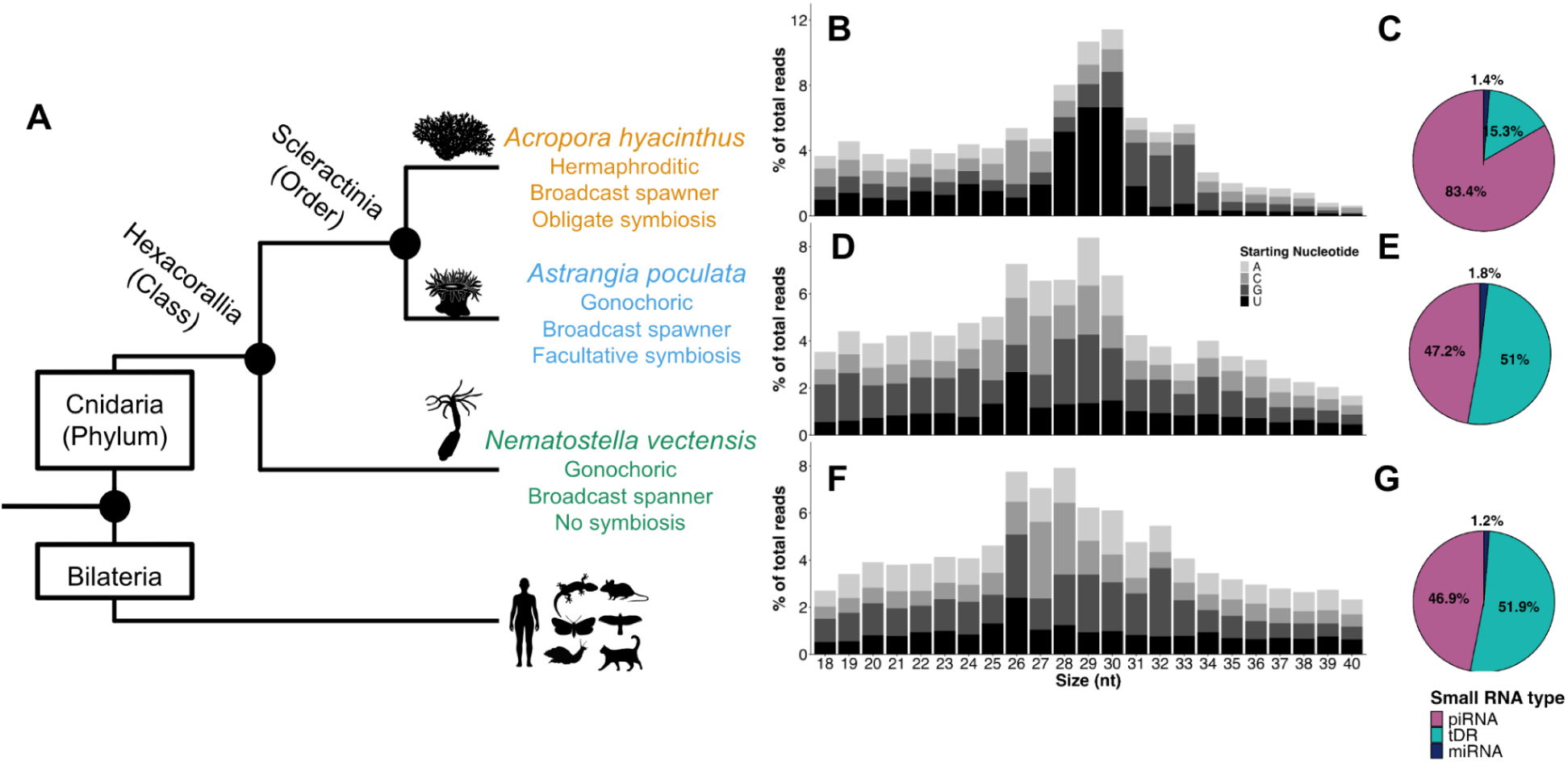
Phylogenetic relationships and small RNA profiles of three cnidarian species. A) Simplified cladogram illustrating the phylogenetic relationships among the three cnidarian species examined in this study and the bilaterian outgroup. Reproductive and symbiotic life-history characteristics are listed for each species. Silhouette images were obtained from https://www.phylopic.org/. B, D, F) Size distribution (18–40 nt) 5’ starting nucleotide composition of trimmed small RNA sequencing reads. Data is shown as percentage of total reads for B) *Acropora hyacinthus*, D) *Astrangia poculata*, and F) *Nematostella vectensis*. Relative proportions of small RNA type (piRNAs, tDRs, and miRNAs) examined in this study in C) *Acropora hyacinthus*, E) *Astrangia poculata*, and G) *Nematostella vectensis*. Percentages indicate the relative abundances of each small RNA type. Alt-text: Small RNA profiles vary among three cnidarian species, with *Acropora hyacinthus* showing piRNA dominance and *Astrangia poculata* and *Nematostella vectensis* exhibiting more balanced piRNA and tRNA-derived RNA proportions.

For *Nematostella vectensis* (gonochoric broadcast spawners; Figure 1A), spawning was induced and sperm samples were collected from individual males as described in Glass et al. (2025).

Briefly, adult males from a laboratory line in culture for ∼2 yr and sexed through previous spawning observations were individually placed in plastic cups of 0.2 µm filtered artificial seawater (ASW; ∼25 mL; 12 psu), and spawning was induced by exposing animals to bright lights and elevated temperatures (∼25°C vs. 18°C ambient culture temperature) for 14 h. Sperm was concentrated into pellets via repeated centrifugation as described above, and pellets were stored at –80°C until further processing.

For *Acropora hyacinthus* (hermaphroditic broadcast spawners; Figure 1A), spawning was observed on November 9, 2023, eleven days after the full moon. Adult colonies (20–30 cm in diameter) were collected from the north shore lagoon in Mo’orea, French Polynesia (–17.48681, –149.88781) on November 3 and 4, transported back to Gump Station in coolers of seawater, and placed in flow-through tanks (∼3000 L) with ambient seawater. On November 9, colonies released egg-sperm bundles, which were collected with 153 µm mesh-bottom containers. Next, 5 mL of bundles were added to a 50 mL falcon tube and 45 mL of FSW was added. Tubes were then laid on their sides and periodically rocked until bundles dissociated. Following bundle disassociation, sperm was collected and concentrated into pellets as described above. Pellets were stored at –80°C until further processing.

### Total RNA isolation

Sperm lysis buffer (33.3 µL; 6.4 M Guanidine HCl, 5% Tween 20, 5% Triton, 120 mM EDTA, and 120 mM Tris pH 8.0), proteinase K (3.3 µL; 20 mg/mL), and 1 M dithiothreitol (3.3 µL) were added to each sample and mixed until the sperm pellet was broken up. Samples were incubated with gentle shaking for 15 min at 60°C. Following incubation, 100 µL of nuclease-free water was added to each sample and mixed well (pipette, vortex, spin down) to ensure the sample was resuspended.

A phenol-chloroform extraction method was used to extract RNA. First, Phase Lock Gel Heavy 2 mL tubes (QuantaBio: 2302830) were centrifuged at 14,000 rpm for 1 min at 4°C. Each sample was then added to a prespun Phase Lock tube, and 50 µL of 1-bromo-2 chloropropane (BCP; Molecular Research Center, Inc; BP151) was added before inverting the tubes at least 10x to mix thoroughly. Samples were then centrifuged at 14,000 rpm for 4 min at 4°C, and the aqueous layer of the resulting phase separation was pipetted into a low-adhesion tube. Next, 110 µL of ice-cold isopropanol and 1 µL of GlycoBlue (Invitrogen, AM9515) was added to each tube. Tubes were vortexed to mix and placed at –20°C for RNA to precipitate for at least 1 h. Following precipitation, tubes were centrifuged at 14,000 rpm for 15 min at 4°C. The supernatant was then discarded and the RNA pellet was washed with 1 mL of ice-cold 70% ethanol. Tubes were centrifuged again at 14,000 rpm for 15 min at 4°C, and all ethanol was removed. The RNA pellet was reconstituted in 8 µL of nuclease-free water.

### Small RNA isolation

Small RNAs were isolated from total RNA by gel extraction based on nucleotide length. First, 8 µL of Invitrogen Gel-Loading Buffer II (ThermoFisher Scientific: CAT #AM8546G) was added to the reconstituted RNA, which was then run on a denaturing gel (15% polyacrylamide, 7 M urea) at 14 W for 30 min. The gel was then stained with 1 μL SYBR Gold Nucleic Acid Gel Stain (ThermoFisher Scientific: CAT #S11494) in ∼100 mL 1x TBE and washed in ∼100 mL 1x TBE, each for 5 min with gentle rocking. A razor blade was then used to excise sections for each lane containing 18–40 bp RNAs as indicated by a custom RNA ladder run on each gel.

Individual gel slices were placed in 1.5 mL low-adhesion microcentrifuge tubes, crushed with a disposable pestle, and 700 µL of gel elution buffer (10 mM Tris-HCl pH 7.5, 1 mM EDTA, 300 mM NaCl) was added to each for overnight extraction at room temperature with shaking (1000 rpm). The following day, buffer containing eluted smRNAs was transferred to a 0.45 μm Costar® Spin-X® centrifuge tube filter (CORNING: CAT # 8170), which was then centrifuged at 10,000 rpm for 1 min at 4°C. The eluate was transferred to a fresh 1.5 mL low-adhesion microcentrifuge tube and 1 μL of GlycoBlue was added, followed by isopropanol precipitation and washing with ethanol as described above.

### Library preparation and sequencing

Washed small RNA pellets were reconstituted in a solution containing RNA 5’ Pyrophosphohydrolase (RppH; New England Biolabs M0356S) prepared according to the manufacturer’s instructions, and then incubated at 37°C for 1 h. Next, RppH-treated smRNAs were again precipitated using isopropanol before library preparation using the Tru-Seq small RNA protocol (Illumina RS-200). Briefly, smRNAs were ligated to 5’ and 3’ adapters, reverse transcribed to cDNA, amplified via PCR, and gel purified (8% TBE polyacrylamide). Libraries were then pooled at equimolar concentration and the pooled library was sequenced using the NextSeq 1000 platform (Illumina) generating 75-bp single-end reads.

### Genomic resources

Genomes, protein and gene sequences, and annotation files were downloaded from the following sources. *Acropora hyacinthus* data was downloaded from GCA_020536085.1 and https://drive.google.com/drive/u/1/folders/15dh6eE8q850frTRVtglndXtzxSwj673g, with an estimated genome size of 446 Mbp, N50 of 26.53 Mbp, and 27,110 protein coding genes (López-Nandam et al. 2023). *Astrangia poculata* data was downloaded from https://zenodo.org/records/14110456, with an estimated genome size of 458 Mbp, N50 of 31 Mbp, and 38,998 protein coding genes (Stankiewicz et al. 2025). *Nematostella vectensis* genome and genomic resources were downloaded from https://simrbase.stowers.org/starletseaanemone, with an estimated genome size of 244 Mb, N50 of 2.1 Mbp, and 24,525 protein coding genes (Zimmermann et al. 2023). To ensure consistent functional annotation across species, protein sequences were queried against the UniProtKB/Swiss-Prot database (accessed on August 25, 2026) using blastp v2.15.0 (Altschul et al. 1990). Output provided protein names and associated gene ontology (GO) terms for downstream comparative analyses. Versions of the genomic resources utilized in this study have been archived on Zenodo (Ashey and Barott 2026).

### Read processing

Fastp v0.23.2 (S. Chen et al. 2018) was used to trim raw reads and low quality sequences and read length limits were set to 15–50 bp. Trimmed read quality was assessed with FastQC v0.12.1 (Andrews 2010) and MultiQC v1.12 (Ewels et al. 2016).

### MicroRNA analysis

To identify novel and known miRNAs, ShortStack v4.1.2 (Axtell 2013; Shahid and Axtell 2014) was used with the parameter –dn_mirna, which identifies putative novel miRNAs by searching read-genome alignments for regions of high coverage and evaluating neighboring regions based on a set of miRNA precursor characteristics. A database of known cnidarian mature miRNAs (miRBase and cnidarian miRNA sequences from previously published studies) was used as a reference (Ashey, Rodriguez-Casariego, et al. 2026). ShortStack results were then filtered for novel miRNAs and known miRNAs identified, retaining those with ≥5 counts across all replicates. miRNA-mRNA interactions were evaluated in both the mRNA coding sequence (CDS) and the 3’ untranslated region (3’UTR), as cnidarian miRNAs can bind to both regions (Moran et al. 2014; Praher et al. 2021). Coding sequences were identified in the species’ genome feature files, which are archived on Zenodo (Ashey and Barott 2026). The 3’UTRs were not annotated in these genomes; therefore, the 1000 bp on the right flank of an mRNA was classified as the 3’UTR (Ashey, Rodriguez-Casariego, et al. 2026). If a 3’UTR was overlapping a gene, the overlapping portions of the 3’UTR were subtracted using bedtools v2.31.1 (Quinlan and Hall 2010). MiRanda v3.3 was used to predict miRNA gene targets with strict seed binding invoked (Enright et al. 2003). To minimize false positives, an energy threshold of ΔG_duplex_ ≤ –20 kcal/mol, a minimum alignment score of 140, and at least 13-bp binding between miRNA and their putative targets were required. The proportion of targets corresponding to 3’UTRs (“3’ UTR only”), CDS regions (“CDS only”), or both (“Both”) was calculated.

### PiwiRNA analysis

Small RNA reads were filtered to keep reads between 25–35 bp and to remove low complexity reads using scripts (TBr2_collapse.pl; TBr2_length-filter.pl; TBr2_duster.pl) from the NGS toolbox (Rosenkranz et al. 2015). To remove non-piRNA reads, short reads were mapped to miRNA, tRNA and rRNA sequences with SortMeRNA v4.3.6 (Kopylova et al. 2012). Reference tRNA and rRNA sequences were generated de-novo using tRNAscan-se v2.0.12 (Chan et al. 2021) and barrnap v0.9 (Seemann 2013), respectively. miRNA sequences were generated from ShortStack, as described above.

Sequences that did not align to the known small RNA sequences were mapped to their respective species reference genome using sRNAmapper.pl from NGS toolbox with the flag –alignments best (Rosenkranz et al. 2015). Origin assessment for multi-mapping reads was done using reallocate.pl from the NGS toolbox (Rosenkranz et al. 2015). The resulting sequences were considered putative piRNAs if they were present in at least three replicates with a minimum overlap of 50% of the piRNA length. Ping-pong amplification signatures were evaluated using PPmeter v0.4 (Jehn et al. 2018). Secondary piRNA processing via reciprocal cleavage was identified by calculating the probability of a 10-nucleotide 5’ overlap between sense and antisense piRNAs, with statistical significance determined by Z-scores > 1.96 (p<0.05) (Jehn et al. 2018).

piRNA overlap with repetitive and genomic coordinates were characterized by intersecting piRNA coordinates with annotated repetitive and genomic elements using bedtools intersect v2.31.1 (Quinlan and Hall 2010). Repetitive elements (i.e., transposable elements or TEs) in the reference genomes were annotated with RepeatModeler v2.0.6 (Flynn et al. 2020) and RepeatMasker v4.15.5 (Smit et al. 2015). Genomic elements were obtained by each species’ genome feature files, and classified into overlaps of “TEs only”, “Genes only”, “Both” (overlapping both a gene and TE), or “Intergenic” (no overlap with genes or TEs).

### tRNA-derived RNAs analysis

To identify tRNA-derived RNAs (tDRs), trimmed small RNA reads were aligned against species-specific tRNA sequences predicted as described above by tRNAscan-se. Alignment was performed with bowtie2 v2.5.2 (Langmead and Salzberg 2012) with strict parameters (--very-sensitive-local –N 1, –L 12, –end-to-end) to prioritize high confidence alignments. To classify mapped tDRs into structural types, a custom Python script evaluated read start and end positions relative to the full-length reference tRNA. Reads were categorized into three distinct types: 5’ tDRs (alignment start coordinates within ≤3 bp of the 5’ terminus), 3’ tDRs (alignment end coordinates within ≤3 bp of the 3’ terminus), and internal tDRs (i-tDRs; mapping internally within the tRNA body without overlapping either terminal boundary). A ≤3 bp terminal window was chosen to account for flexibility in endonucleolytic cleavage and minor 5’ or 3’ terminus trimming during tDR biogenesis.

To evaluate tDR spatial coverage profiles across mature tRNAs, positional read depth of the tDRs was calculated across full-length reference tRNA sequences for each species using custom Python scripts. Positional coverage data was aggregated across individual tRNA genes by amino acid and anticodon isoacceptor families (e.g., GlyGCC, AspGTC, etc). The four most abundant tRNA isoacceptors per species were identified based on normalized reads per million (RPM).

Nucleotide coverage distribution curves were constructed to visualize relative terminal enrichment (i.e., 5’ vs. 3’) across mature tRNA bodies for each species.

### Homology analysis

To determine whether sperm small RNAs target functionally conserved genes across cnidarians, orthologous gene clusters (orthogroups) were identified across *A. hyacinthus, A. poculata,* and *N. vectensis* using Broccoli v1.2 (Eklund et al. 2014). Orthogroups were evaluated to determine whether they contained genes targeted by miRNAs or genes overlapping with predicted piRNAs. Shared orthogroups containing targets or overlaps across all three species were extracted to identify conserved paternal small RNA targets.

To determine conserved biological roles, functional gene ontology (GO) enrichment was conducted on the shared target genes for miRNAs and the shared gene overlaps for piRNAs using the R package topGO v2.60.1 (Alexa and Rahnenführer 2024). Significant enrichment was assessed for Biological Processes (BP) and Molecular Functions (MF) using Fisher’s exact test paired with the weight01 algorithm from topGO (significance threshold of p<0.05). For each species and small RNA class (miRNA gene targets and piRNA-overlapping genes), the top 100 enriched GO terms ranked by p-value were retained. Finally, to identify core genes underlying these conserved processes, we cross-referenced the genes with significant GO terms across all three species to identify shared candidate genes that were both orthologous and associated with the shared enriched functional term. Examining the specific gene annotations (i.e., not the GO term but the actual name of the gene from Swissprot) enabled us to move beyond broad functional categories and resolve specific pathways putatively targeted by paternal small RNAs. Functional analysis was done using R v4.5.0 (R Core Team 2024).

## Results

### Small RNA composition

Small RNA sequencing of the sperm of three cnidarian species revealed distinct size distributions and RNA type compositions (Figure 1). *Acropora hyacinthus* exhibited a pronounced peak at approximately 28–29 nt (Figure 1B), whereas *Astrangia poculata* and *Nematostella vectensis* displayed broader distributions, with elevated read abundance across approximately 26–30 nt (Figure 1D,F).

The relative abundance of small RNA types also differed among species. *A. hyacinthus* sperm was predominantly composed of piRNAs (83.4%; Figure 1C), while *A. poculata* and *N. vectensis* exhibited more balanced proportions of piRNAs and tDRs (Figure 1E,G). miRNAs represented a comparatively smaller fraction of the annotated small RNA population in all three species.

### MicroRNAs

MicroRNAs were successfully detected in the sperm of all three cnidarian species by mapping small RNA sequencing reads against previously established species-level repertoires (*Acropora hyacinthus*: (Gajigan and Conaco 2017; Praher et al. 2021; Ashey, Rodriguez-Casariego, et al. 2026); *Astrangia poculata*: (Ashey, Gilligan, et al. 2026); *Nematostella vectensis*: (Moran et al. 2014)). In total, 25, 7, and 12 distinct miRNAs were identified in *A. hyacinthus, A. poculata,* and *N. vectensis* sperm, respectively, all of which corresponded to known loci with no novel miRNAs detected (Table S1). Notably, miR-100 was the sole conserved miRNA identified in sperm across all three species.

To evaluate the functional potential of sperm miRNAs, target prediction was performed using the coding sequence (CDS) and 3’ untranslated regions (3’UTRs) of transcripts as potential targets. This analysis revealed a high degree of sequence complementarity between the identified miRNAs and their putative targets across all taxa (Figure 2A). In *A. hyacinthus*, 6,204 unique mRNAs were identified as putative targets, with 97.66% exhibiting a binding complementarity greater than 8 base pairs (bp), the canonical seed sequence length typical of bilaterian miRNAs. Binding interactions in *A. hyacinthus* peaked at 19 bp of shared complementarity, accounting for 33.31% of all predicted interactions. Similarly, 3,294 unique mRNA targets were predicted in *A. poculata*, with 98.24% displaying high-affinity binding (>8 bp) and a peak binding frequency at 20 bp (32.05%). In *N. vectensis*, 6,624 unique mRNAs were identified as targets (97.96% with >8 bp binding), with the highest binding frequency occurring at 18 bp (22.79%).

**Figure 2.**
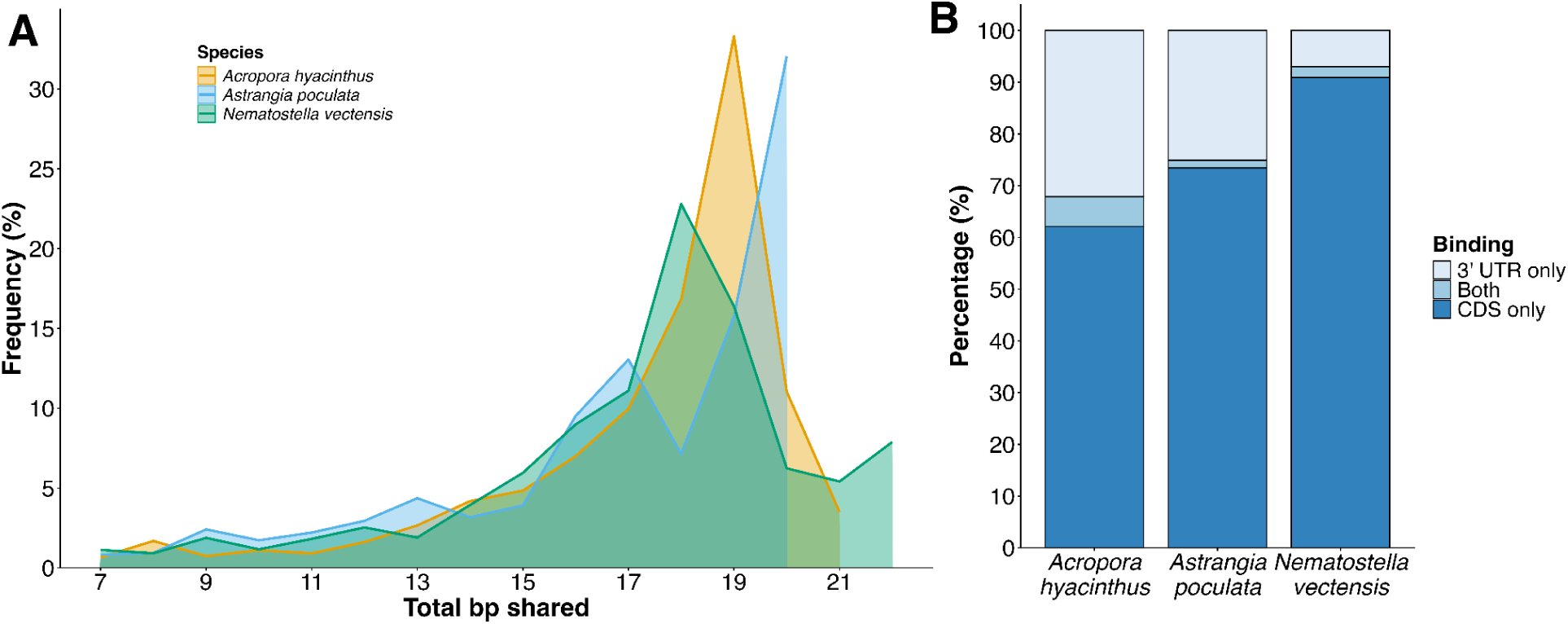
Characteristics of predicted microRNA targeting across three cnidarian species. A) Frequency distribution of total number of complementary base pairs (bp) shared between mature miRNAs and their putative mRNA targets in *Acropora hyacinthus* (orange), *Astrangia poculata* (blue), and *Nematostella vectensis* (green). B) Proportion (%) of predicted miRNA binding sites exclusively within coding sequences (“CDS only”), exclusively within the 3’ untranslated region (“3’ UTR only”), or spanning both regions (“Both”) across the three species. Alt-text: Predicted miRNA–target interactions show species-specific distributions of sequence complementarity, with all three cnidarians exhibiting frequent interactions involving 18–20 complementary base pairs.

While the distribution of predicted miRNA binding sites across target transcripts varied by species, binding sites were consistently enriched within coding sequences (CDS) across all three species (Figure 2B). The majority of putative targets contained binding sites localized exclusively to the CDS region, ranging from 62.11% in *A. hyacinthus* and 73.44% in *A. poculata* to a high of 90.92% in *N. vectensis* (Figure 2B). Conversely, putative miRNA targets containing predicted binding in the 3’UTR were less frequent, particularly in *N. vectensis,* where only 6.98% of targets were associated with the 3’ UTR (Figure 2b). A comparatively higher proportion of 3’ UTR-contained targets was observed in the scleractinian corals, representing 25.08% of targets in *A. poculata* and 32.10% in *A. hyacinthus* (Figure 2B). Targets containing predicted binding sites across both regions constituted a minor fraction of the total profile in all species (Figure 2B).

To determine whether sperm miRNAs putatively regulate conserved functions across cnidarians, we evaluated orthogroups containing orthologous mRNA targets in *A. hyacinthus*, *A. poculata*, and *N. vectensis* (Figure 3). Comparative orthology analysis revealed 1,535 orthogroups that contained mRNA targets from the three species (Figure 3A). Functional enrichment analysis of these shared targets identified 21 significantly enriched GO terms across the species (Table S2). Cross-referencing these genes with Swissprot annotation revealed 36 shared candidate genes that were both orthologous across the three species and associated with the shared enriched GO terms. These conserved targets were related to post-transcriptional and epigenetic regulation, cell adhesion dynamics, and signal transduction (Figure 3B).

**Figure 3.**
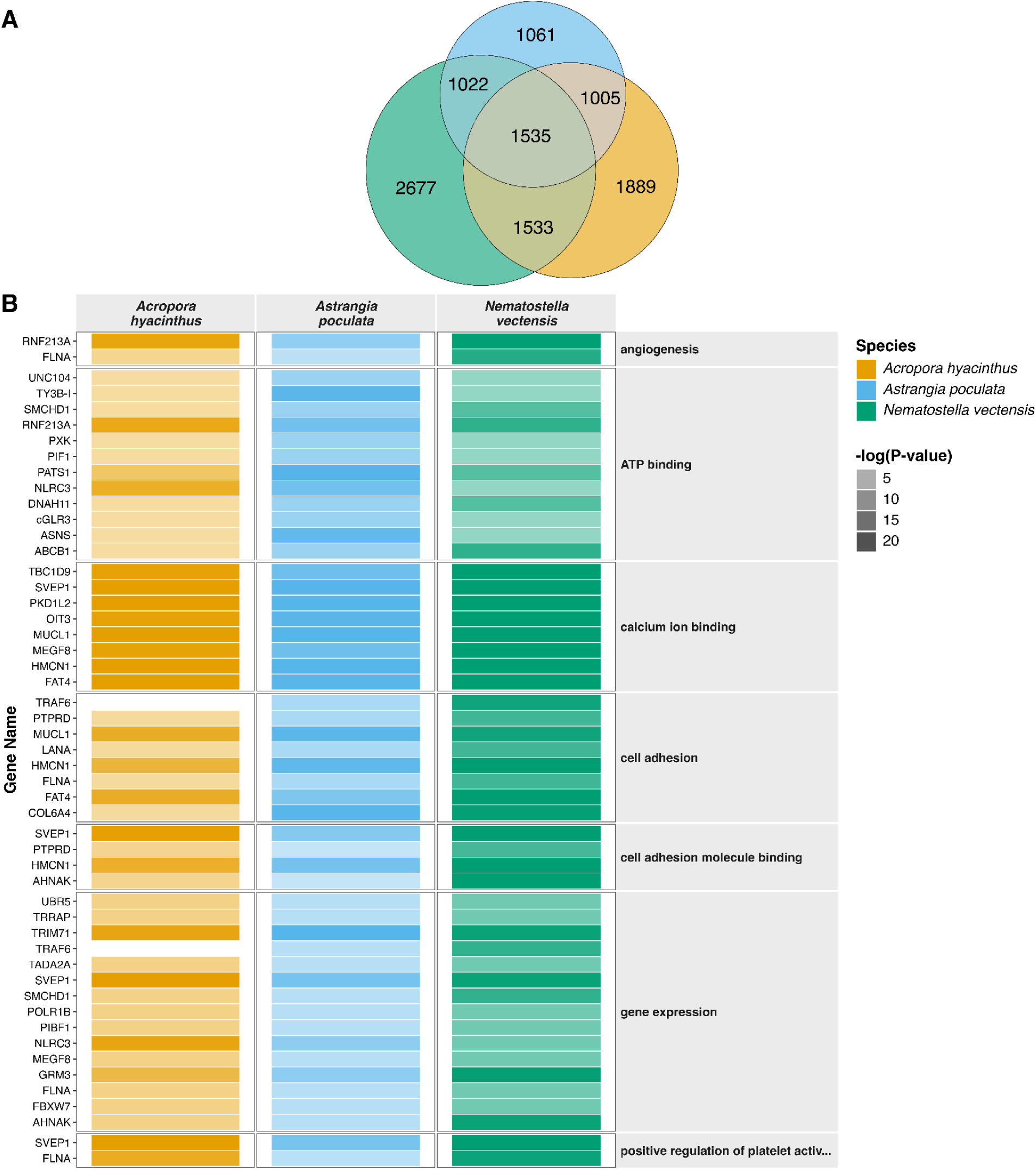
Functional conservation of putative miRNA targets across cnidarians sperm. A) Venn diagram displaying the overlap of orthogroups with predicted targets across *Acropora hyacinthus* (orange)*, Astrangia poculata* (blue), and *Nematostella vectensis* (green). A total of 1,535 orthogroups were shared among all three species. B) Heatmap showing predicted miRNA target genes associated with significantly enriched gene ontology (GO) terms shared across the three species. Rows represent individual target genes, grouped by associated GO terms. Color opacity represents enrichment significance, expressed as –log(P-value) derived from Fisher’s exact test. Only GO terms associated with more than one gene were included for visualization. Complete enrichment results are provided in Table S2. Alt-text: Predicted miRNA targets include 1,535 shared orthogroups across three cnidarian species, with conserved gene associations spanning multiple functional categories.

Post-transcriptional and epigenetic regulation was reflected by candidate annotated genes under the shared enriched GO terms gene expression and ATP binding (Figure 3B). Shared candidate genes underlying these terms included transcriptional and chromatin remodeling regulators, such as transformation/transcription domain-associated protein (TRRAP), transcriptional adapter 2-alpha (TADA2A), DNA-directed RNA polymerase I subunit RPA2 (POLR1B), structural maintenance of chromosomes flexible hinge domain containing 1 (SMCHD1), Transposon Ty3-I Gag-Pol polyprotein (TY3B-I), and F-box/WD repeat-containing protein 7 (FBXW7).

Post-transcriptional protein regulation was represented by E3 ubiquitin-protein ligases (UBR5 and TRIM71). These GO terms also included shared genes related to signaling, including ABC transporter B family member 1 (ABCB1), Cyclic GMP-AMP synthase-like receptor 3 (cGLR3), serine/threonine-protein kinase (PATS1), PX domain-containing protein kinase-like protein (PXK), Metabotropic glutamate receptor 3 (GRM3), and TNF receptor-associated factor 6 (TRAF6; Table S2).

Cellular architecture was implicated under the shared enriched GO terms cell adhesion, cell adhesion molecule binding, and angiogenesis (Figure 3B). Shared candidate genes annotated with these structural terms included neuroblast differentiation-associated protein (AHNAK), filamin-A (FLNA); Hemicentin-1 (HMCN1); Protocadherin Fat 4 (FAT4); Laminin subunit alpha (LANA); Collagen alpha-4(VI) chain (COL6A4); Polycystin-1-like protein 2 (PKD1L2); Sushi, von Willebrand factor type A, EGF and pentraxin domain-containing protein 1 (SVEP1); Receptor-type tyrosine-protein phosphatase delta (PTPRD); and Mucin-like protein (MUCL1; Table S2).

### PiwiRNAs

The presence of sperm piRNAs was evaluated by assessing ping-pong processing signatures across identified piRNA transcripts. In the canonical ping-pong amplification loop, secondary biogenesis is driven by reciprocal cleavage events that generate a characteristic 10-nucleotide 5’ overlap between sense and antisense piRNAs (Siomi et al. 2011). Across *A. hyacinthus, A. poculata,* and *N. vectensis*, piRNAs displayed this characteristic 10-nucleotide 5’ overlap signature (Figure 4A–C; Z-scores = 8.31, 11.80, and 5.42, respectively), confirming secondary piRNA processing via the ping-pong amplification loop.

**Figure 4.**
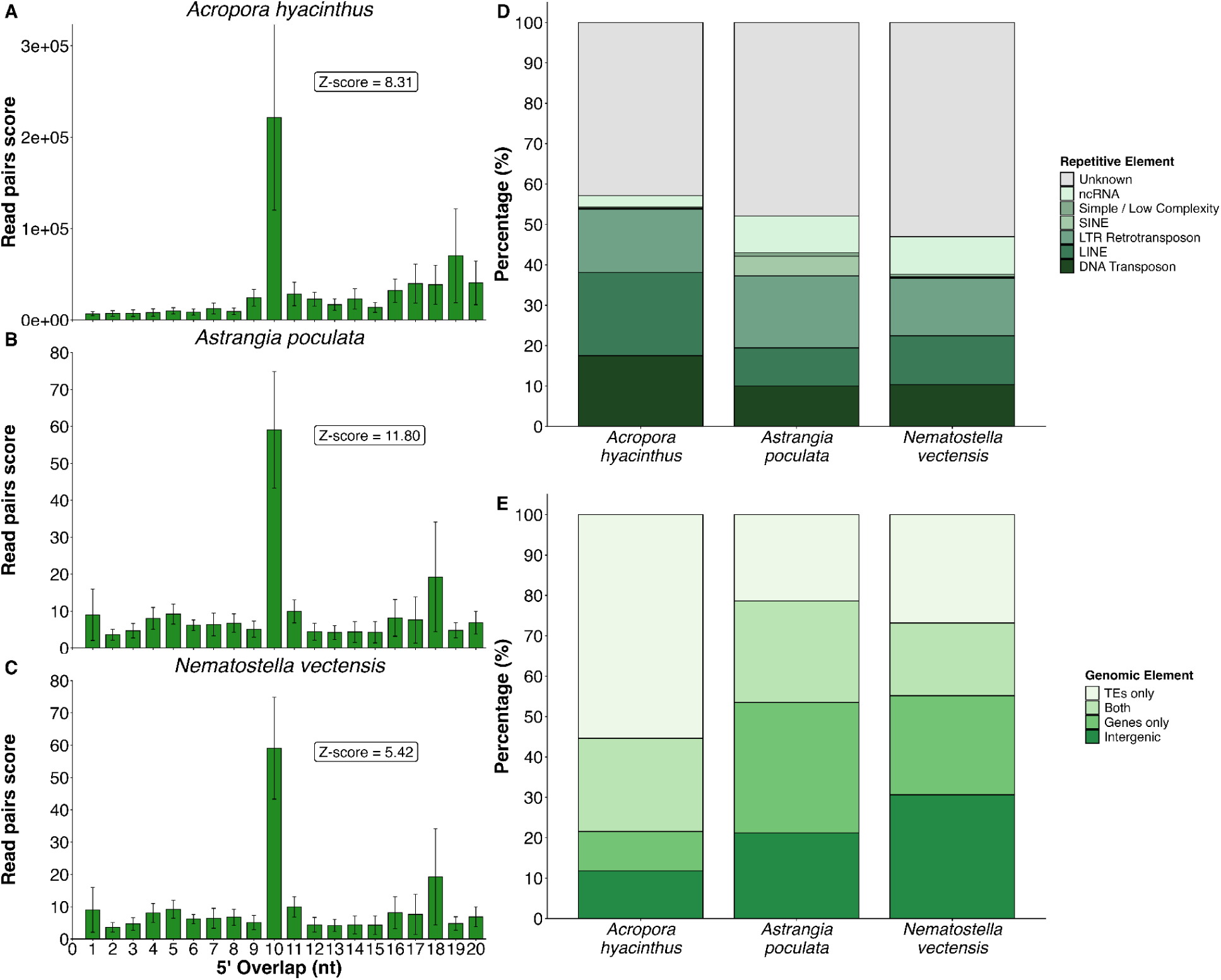
piRNA processing signatures and genomic distribution of sperm piRNAs across three cnidarian species. A–C) Ping-pong amplification signatures determined by 5’ overlap probability between sense and antisense piRNAs in A) *Acropora hyacinthus*, B) *Astrangia poculata*, and C) *Nematostella vectensis*. The peak at 10 nucleotides (nt) indicates the characteristic 10-nt overlap associated with secondary piRNA biogenesis through the ping-pong amplification cycle. Z-scores are shown for each species. D) Relative proportion (%) of sperm piRNAs overlapping with specific repetitive elements across species, including DNA transposons, long interspersed nuclear elements (LINEs), long terminal repeat retrotransposons (LTR retrotransposons), short interspersed nuclear elements (SINEs), simple / low complexity repeats, non-coding RNAs (ncRNAs), and unknown repetitive sequences. E) Genomic distribution (%) of sperm piRNAs overlapping with annotated genomic elements. Genomic elements included transposable elements only (“TEs only”), genes only (“Genes only”), both genes and TEs (“Both”), and intergenic regions (“Intergenic”). Alt-text: All three cnidarian species exhibit a pronounced ten-nucleotide piRNA overlap signature, with species-specific distributions across repetitive elements and genomic regions.

Sperm piRNAs in all three species overlapped with diverse repetitive elements, including DNA transposons, long interspersed nuclear elements (LINEs), long terminal repeat retrotransposons (LTR retrotransposons), short interspersed nuclear elements (SINEs), simple/low complexity repeats, non-coding RNAs (ncRNAs), and unknown repetitive sequences (Figure 4D).

Unannotated repeats represented the largest category of repeat-associated piRNAs in each species: 42.87% were unknown in *A. hyacinthus*, 47.90% in *A. poculata*, and 53.03% in *N. vectensis* (Figure 4D). High proportions of unknown repeat elements are characteristic of cnidarian genome annotations due to divergent repetitive structures (Noel et al. 2023; Conn et al. 2025; Fiesinger et al. 2026). Among annotated repetitive elements, piRNAs most frequently overlapped with LINEs in *A. hyacinthus* (20.60%) and LTR retrotransposons in *A. poculata* (17.83%) and *N. vectensis* (14.26%) (Figure 4D).

Examining piRNA overlaps with general genomic elements revealed that sperm piRNAs overlap with both transposable elements (TEs) and genes (Figure 4E). Total TE-associated piRNAs (encompassing “TEs only” and “Both”) accounted for 78.42% of the piRNAs in *A. hyacinthus*, 46.51% in *A. poculata*, and 44.81% in *N. vectensis* (Figure 4E). A substantial fraction of piRNAs in all species also overlapped with genes (encompassing “Genes only” and “Both”), accounting for 32.83% of piRNAs in *A. hyacinthus*, 57.41% in *A. poculata*, and 42.54% in *N. vectensis* (Figure 4E).

To assess whether piRNA-overlapping genes overlap with conserved genes, we evaluated orthogroups containing genes overlapping piRNAs in *A. hyacinthus, A. poculata,* and *N. vectensis*. There were 1614 piRNA-overlapping orthogroups across the three species (Figure 5A). Functional enrichment of these shared orthogroups identified 38 significantly enriched shared GO terms across species (Table S3). Cross-referencing these annotated genes revealed 56 candidate genes that were both orthologous across all three species and associated with 34 of the shared enriched GO terms (Table S3). These conserved piRNA-overlapping genes were consistently enriched with terms related to translational and ribosomal machinery, nucleic acid dynamics and genomic maintenance, and cytoskeletal organization and transport.

**Figure 5.**
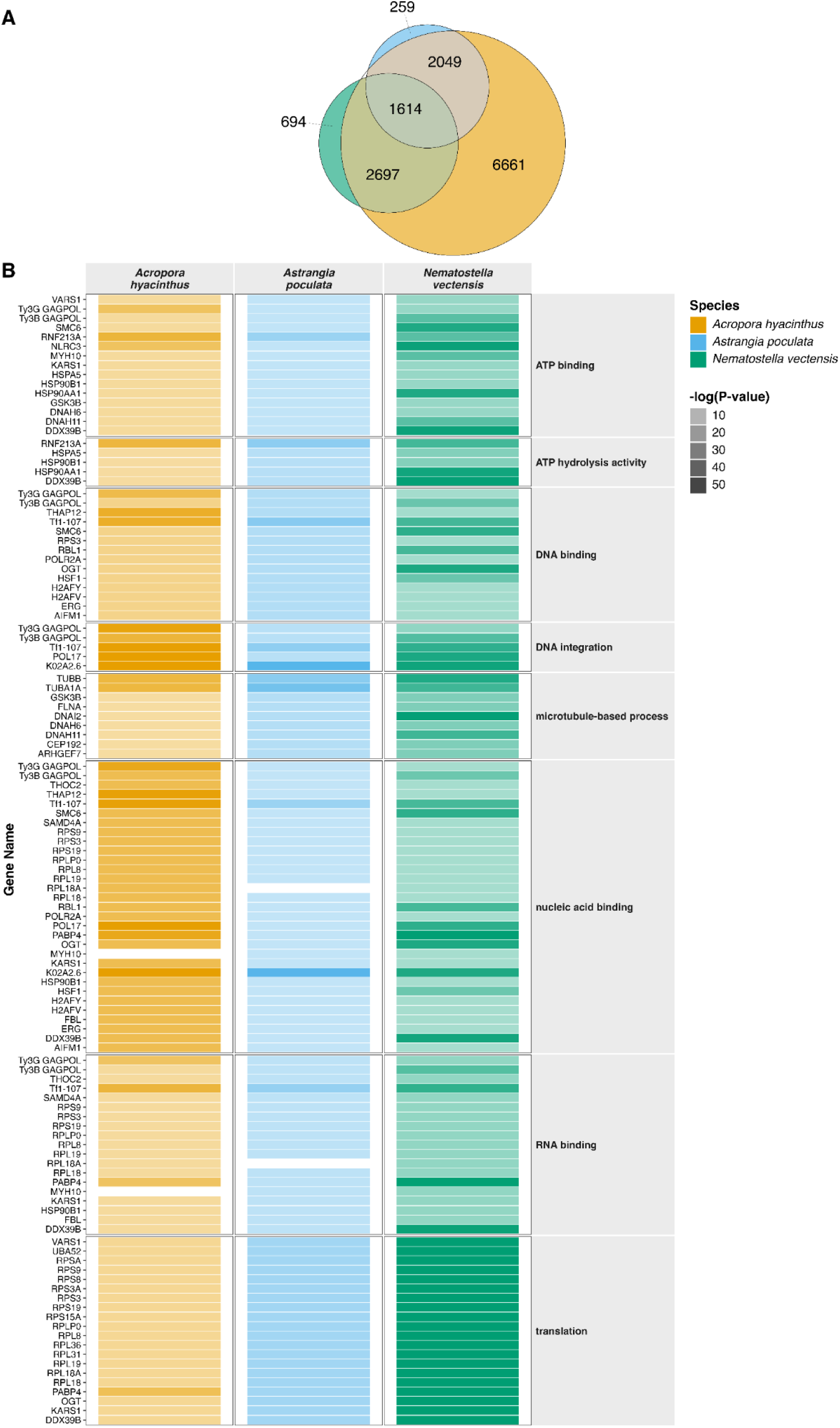
Functional conservation of piRNA-associated overlaps across cnidarian sperm. A) Venn diagram displaying the overlap of orthogroups containing genes associated with piRNA genomic overlaps among *Acropora hyacinthus* (orange)*, Astrangia poculata* (blue), and *Nematostella vectensis* (green). A total of 1,614 orthogroups were shared across all three species. B) Heatmap showing gene overlaps associated with significantly enriched gene ontology (GO) terms shared across the species. Each row corresponds to an individual gene, clustered by its GO term. Color opacity represents enrichment significance, expressed as –log(P-value) derived from Fisher’s exact test. Only GO terms associated with more than one gene were included for visualization. Complete enrichment results are provided in Table S3. Alt-text: Three cnidarian species share 1,614 piRNA-associated orthogroups, with overlapping genes enriched in functions related to nucleic acid binding, RNA binding, and translation.

Shared enriched GO terms such as RNA binding and translation indicate potential regulation of the genes overlapping with piRNAs across the three species (Figure 5B). Genes encoding structural components of both the large (RPL) and small (RPS) ribosomal subunits were consistently identified as piRNA-overlapping genes, alongside aminoacyl-tRNA synthetases including lysine–tRNA ligase (KARS1) and valine–tRNA ligase (VARS1; Table S3).

Nucleic acid regulatory functions were represented under the shared enriched GO terms ATP binding, DNA binding, DNA integration, and nucleic acid binding (Figure 5B).

piRNA-overlapping genes in these enriched terms included transcriptional and post-transcriptional regulators, such as transcriptional regulator ERG (ERG), DNA-directed RNA polymerase II subunit RPB1 (POLR2A), sterile alpha motif domain containing 4A (SAMD4A), polyadenylate-binding protein 4 (PABP4), rRNA 2’-O-methyltransferase fibrillarin (FBL), spliceosome RNA helicase (DDX39B), and heat shock factor protein 1 (HSF1). Histone variants, including H2AFV and H2AFY, and structural maintenance of chromosome protein 6 (SMC6) were also found in the enriched terms. Additionally, transposon genes were present in the enriched terms, such as Tf1-107, Ty3B-GAGPOL, and Ty3G-GAGPOL (Table S3).

Cytoskeletal structure and intracellular transport functions were found in the enriched GO term microtubule-based process. Shared candidate genes in this enriched term included centrosomal protein of 192 kDa (CEP192), dynein axonemal heavy chain 6 and 11 (DNAH6 and DNAH11), dynein intermediate chain 2 (DNAI2), filamin-A (FLNA), tubulin alpha-1 chain (TUBA1A), tubulin beta chain (TUBB), and Rho guanine nucleotide exchange factor 7 (ARHGEF7; Table S3).

### tRNA-derived RNAs

In all species, 5’ tDRs comprised the majority of mapped fragments, accounting for 52.93% of mapped tDRs in *A. hyacinthus*, 57.44% in *A. poculata*, and 46.55% in *N. vectensis* (Figure 6A). Internal tDRs (i-DRs) comprised the second most abundant type, accounting for 39.06% in *A. hyacinthus*, 29.69% in *A. poculata*, and 30.82% in *N. vectensis* (Figure 6). Across all species, the 3’ tDRs represented the smallest fraction, with 8.02% of tRNA reads in *A. hyacinthus*, 12.87% in *A. poculata*, and 22.64% in *N. vectensis* (Figure 6A). These fragments represent downstream products of nucleolytic tRNA cleavage, typically initiated at the anticodon loop (CITE).

**Figure 6.**
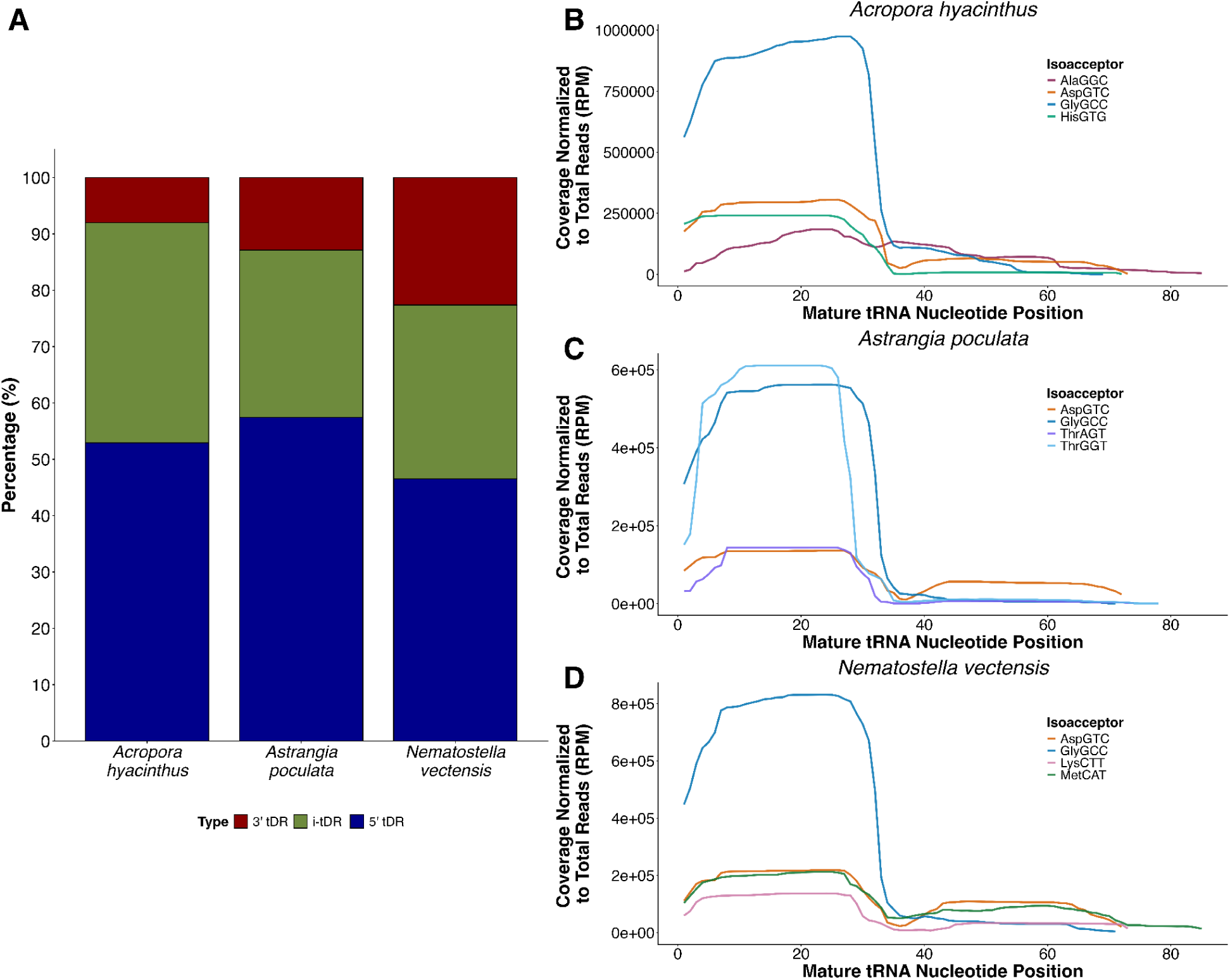
Structural origins and positional coverage of tDRs across cnidarian sperm. A) Proportion of mapped tDRs classified by their position of origin within mature tRNA sequences: five-prime tDRs (5′ tDRs; blue), internal tDRs (i-tDRs; green), and three-prime tDRs (3′ tDRs; red). Five-prime and three-prime tDRs were defined as reads with alignment start or end positions, respectively, within three nucleotides of the corresponding tRNA terminus. Internal tDRs mapped within the tRNA sequence without overlapping either terminal region. B–D) Single-nucleotide positional coverage of reads mapped to mature tRNA sequences, normalized as reads per million (RPM) of total reads to the four most abundant tRNA isoacceptors in B) *Acropora hyacinthus*, C) *Astrangia poculata*, and D) *Nematostella vectensis*. Line colors indicate specific tRNA isoacceptor families as denoted in the species legends. Alt-text: All three cnidarian species show predominantly five-prime and internal tRNA-derived RNAs, with distinct nucleotide coverage profiles across individual tRNA isoacceptors.

However, these relative proportions should be interpreted within the context of ligation-dependent library preparation. Because standard adapters require canonical terminal ends (5′-monophosphates and 3′-hydroxyls), tDRs with non-canonical or modified ends may be underrepresented.

Evaluation of positional coverage across mature tRNAs revealed terminal biases among highly abundant tRNA isoacceptors (Figure 6B–D). For the four most abundant isoacceptor families in each species, coverage was concentrated along the 5’ terminal region (positions ∼1–35 bp).

Notably, fragments derived from GlyGCC and AspGTC were consistently among the top expressed tDRs in all three species, displaying prominent 5’-terminal accumulation peaks. Other top isoacceptors included AlaGGC and HisGTG in *A. hyacinthus*, ThrAGT and ThrGGT in *A. poculata*, and LysCTT and MetCAT in *N. vectensis*, all of which displayed similar 5’ coverage profiles.

## Discussion

### Cnidarian sperm host distinct small RNA types

Our results demonstrate that cnidarian sperm hosts a diverse repertoire of small RNAs including miRNAs, piRNAs, and tDRs. Although all three species contained each of the small RNA types examined, their relative abundance varied. In *A. hyacinthus*, sperm was dominated by piRNAs, whereas *A. poculata* and *N. vectensis* possessed a more balanced proportion of piRNAs and tDRs. Across all three species, miRNAs represented a relatively small proportion of small RNAs profiled. However, low abundance does not necessarily indicate limited regulatory potential.

Recent work by Lee et al. (2026) demonstrated that as few as 200 molecules of sperm-derived miRNAs could induce measurable changes in embryonic gene expression and influence developmental outcomes in mice. Thus, despite their relatively lower abundance in cnidarian sperm, miRNAs may still contribute significantly to paternal regulation of early development.

The substantial representation of tDRs in cnidarian sperm is consistent with observations in model systems, where tDRs constitute a major component of the mature sperm small RNA population and have been implicated in paternal transmission of environmental information (Chen et al. 2016; Sharma et al. 2016; Galambos et al. 2026). In mammalian sperm, piRNAs are typically expressed at higher levels during early spermatogenesis, but as maturation proceeds, piRNAs become less abundant while tDRs become enriched (Sharma et al. 2016; Conine et al. 2018). The relatively higher proportion of piRNAs in mature cnidarian sperm raises questions about if similar small RNA remodeling occurs during cnidarian spermatogenesis. Alternatively, differences in observed abundances may be partly reflected in technical biases associated with RNA extraction and library preparation. Indeed, comparisons of human sperm small RNA populations have demonstrated substantial variation in RNA composition across individuals and populations, potentially reflecting biological, environmental, and technical factors (Mehta and Singh 2025). Together, these findings suggest that sperm small RNA compositions are not uniform across animal lineages and likely reflect a complex interplay of environmental, evolutionary, and technical factors.

### Cnidarian miRNA target features resemble those of plants

Our characterization of miRNA-target interactions revealed regulatory features in cnidarians that parallel plant systems rather than traditional bilaterian models. In bilaterians, miRNAs typically regulate targets through imperfect complementarity, relying on a 6–8 bp “seed sequence” to bind to the 3’UTR of target mRNAs (Axtell et al. 2011). However, in plants, miRNAs tend to bind with near-perfect complementarity to their target (Axtell et al. 2011). Here, we found that miRNAs in our three cnidarian species exhibited high complementarity with their targets, mirroring plant miRNA binding dynamics. This aligns with recent functional evidence from *N. vectensis*, where seed-sequence matches alone were insufficient for miRNA-mediated knockdown, requiring near-full complementary matches to effectively silence their targets (Admoni et al. 2025).

Furthermore, the majority of miRNAs across the three cnidarian species targeted the CDS region rather than 3’UTRs. CDS binding is characteristic of plant miRNA regulatory processes (Ding et al. 2012; Hausser et al. 2013) and has been increasingly identified across cnidarian lineages (Praher et al. 2021). Previous work demonstrated that *N. vectensis* miRNAs in somatic tissues can target different transcript regions (3’ UTR, 5’ UTR, CDS), with a heavy representation in the CDS (Moran et al. 2014). Similarly, most putative mRNA-miRNA binding sites in the cnidarian *Aiptasia* were located in CDS regions (Baumgarten et al. 2018). Computational predictions from corals have also demonstrated that miRNAs are predicted to bind in the CDS region (Despard et al. 2025; Ashey, Gilligan, et al. 2026; Durkin et al. 2026). Collectively, these findings suggest that CDS-directed regulation with high sequence complementarity likely represents a conserved mechanism of miRNA action across cnidarians (Praher et al. 2021).

### Shared miRNA target homologs suggest conserved roles in fertilization and early development related to cellular architecture and transcriptional regulation

The shared miRNA targets identified across *A. hyacinthus, A. poculata,* and *N. vectensis* revealed enriched processes related to post-transcriptional and epigenetic regulation, cell adhesion dynamics, and signal transduction. Because our bioinformatic miRNA target predictions rely on sequence complementarity and energy binding rather than functional knockdowns, these putative interactions warrant future experimental validation. However, they establish a cross-lineage framework for powerful hypothesis generation for the role of sperm small RNAs in cnidarians.

While sperm-borne miRNAs could potentially participate in pre-spawning spermatogenesis functions, their presence in transcriptionally silent sperm strongly points to a post-fertilization role upon delivery to the egg. Indeed, sperm miRNAs in bilaterians are critical reprogrammers of the early embryonic transcriptome (Trigg and Conine 2024; Lee et al. 2026). Therefore, we interpret our putative targets through the lens of their potential functional roles during fertilization and early cleavage, while acknowledging their potential role in spermatogenesis.

The conservation of these targets across ecologically and evolutionary distinct cnidarian lineages suggests an ancient, conserved role for paternal miRNAs in regulating foundational developmental processes. Specifically, the enrichment of putative paternal targets involved in transcriptional machinery (TRRAP, TADA2A, POLR1B) and epigenetic regulation (SMCHD1, UBR5, TRIM71) suggests a mechanism by which paternal miRNAs could actively shape the landscape and timing of early embryonic gene expression. This regulatory timing is particularly critical during the maternal-to-zygotic transition (MZT), a pivotal window where embryonic development depends on the coordinated clearance of maternal transcripts and the initiation of zygotic genome activation (ZGA) (Tadros and Lipshitz 2009; Vastenhouw et al. 2019). Our findings raise the possibility that sperm miRNAs targeting early transcriptional and epigenetic factors may mediate the MZT or early embryogenesis in cnidarians. However, more work is needed to experimentally validate this hypothesis.

TRIM71 was also putatively targeted by sperm miRNAs across the three species. TRIM71 is an E3 ubiquitin ligase, but can also interact with miRNAs and their associated proteins (Chang et al. 2012). Through coordination with miRNAs and Argonaute proteins, TRIM71 promotes embryonic cell proliferation (Chang et al. 2012), while its loss during embryogenesis leads to severe germ cell depletion and developmental arrest (Torres-Fernández et al. 2021). miRNAs putatively targeting their own processing and effector machinery has also been documented in adult cnidarians (Ashey, Rodriguez-Casariego, et al. 2026; Durkin et al. 2026). This suggests that the autoregulatory feedback loop of miRNAs may be a critical feature of cnidarian biology that operates across the entire life cycle.

Terms related to cellular adhesion and architecture were significantly enriched in target orthologs across the three species and suggests paternal miRNAs actively regulate the transition from fertilization to early cleavage/division. In our study, the specific targeting of structural and extracellular matrix transcripts like FLNA, HMCN1, COL6A4, FAT4, and AHNAK indicates that sperm miRNAs may be coordinating cellular division in early development. Early embryonic division is an incredibly sensitive process, requiring precisely timed structural remodeling to ensure proper development (Mu et al. 2022). The putative targeting of these specific transcripts by sperm miRNAs indicate a conserved mechanism by which sperm miRNAs may modulate early embryonic cellular division in cnidarians.

### Sperm piRNAs may serve dual roles in TE and gene regulation

Consistent with the canonical role of piRNAs in maintaining genomic integrity (Czech et al. 2018), we observed considerable sequence overlap between sperm piRNAs and TEs across the three species. However, a substantial fraction of sperm piRNAs also aligned with protein-coding genes. While piRNAs have a canonical function in TE silencing, a growing body of literature illustrates a role for piRNAs in gene regulatory functions (Gebert et al. 2015; Chen et al. 2021). Within cnidarians, gene targeting by piRNAs has been documented in *N. vectensis* (Praher et al. 2017) and *Hydra* (Krishna et al. 2013; Juliano et al. 2014; Teefy et al. 2020). In mammalian systems, functional inhibition of paternal piRNAs from the sperm has negative downstream effects in fertilization success and early development (Wu et al. 2020; Perillo et al. 2023), indicating that sperm piRNAs are not a transcriptional remnant of spermatogenesis but an essential part of development. Our findings suggest that cnidarian sperm piRNAs may simultaneously target TEs and protein-coding genes to impact gene expression regulation. However, experimental validation is needed to fully understand the mechanistic roles of sperm-derived piRNAs in cnidarians.

### Conserved gene-overlapping piRNAs indicate putative paternal control over embryonic translation and transcriptional onset

Because mature sperm are transcriptionally silent, we interpret sperm-borne piRNAs primarily through the lens of post-fertilization early development, while acknowledging their potential prior roles in spermatogenesis (Hong et al. 2025). The consistent overlap of piRNAs expressed in Cnidarian sperm with genes encoding translational machinery points to a potential role in tuning embryonic translation during early development. Across all three species, piRNAs overlapped with genes coding for structural components of the large and small ribosomal subunits (RPL, RPS) and amino-tRNA ligases (KARS1, VARS1). Early metazoan development relies on the selective translation of maternally provisioned mRNAs prior to zygotic genome activation (Tadros and Lipshitz 2009; Vastenhouw et al. 2019). By putatively targeting transcripts associated with protein synthesis, sperm piRNAs delivered at fertilization may modulate translation rates, ribosome assembly, and/or maternal mRNA clearance. In *Drosophila*, piRNA complexes actively contribute to the targeted cleavage and decay of maternal mRNAs in the early embryo (Rouget et al. 2010; Barckmann et al. 2015), and disruption of this piRNA-mediated decay leads to aberrant translational derepression and developmental defects (Rouget et al. 2010). While these studies do not specifically highlight sperm-derived piRNAs, it is plausible that paternal piRNAs delivered from the sperm could similarly contribute to early transcript degradation and translation, providing a mechanism for paternal piRNAs to influence developmental timing in cnidarian embryos.

Developmental timing and zygotic transcriptional onset may be further modulated by sperm piRNAs putatively targeting genes encoding histone variants (H2AFV, H2AFY), chromosome maintenance factors (SMC6), and retrotransposon polyproteins (Tf1-107, GAG-POL variants). Across eukaryotes, piRNAs silence TEs primarily both posttranscriptionally and transcriptionally by remodeling chromatin state. For instance, piRNAs are essential for maintaining repressive H3K9me3 histone modifications on TEs in mammalian germ cells (Pezic et al. 2014) and modulating chromatin accessibility to enforce TE silencing *in vitro* (Iwasaki et al. 2016). The presence of sperm piRNAs overlapping with both architectural genes and retrotransposon sequences in our dataset suggests that cnidarian paternal piRNAs may perform their canonical role of safeguarding genome integrity by suppressing TEs while simultaneously regulating architectural factors to prime the zygotic genome for activation.

### 5’-terminal tDRs identified in three cnidarian species

Here, we report the first identification and structural profiling of tDRs in cnidarians, revealing that tDRs are present in *A. hyacinthus, A. poculata,* and *N. vectensis* sperm, with 5’ tDRs consistently comprising the dominant structural type. In bilaterian models, sperm tDRs accumulate during gamete maturation and serve as key vectors of paternal epigenetic inheritance that shape embryonic early development (Sharma et al. 2016; Conine et al. 2018). For example, in *Caenorhabditis elegans*, sperm tDR accumulation regulates early embryonic gene expression and transmits inherited phenotypes across generations, including starvation resilience and altered heat stress tolerance (Galambos et al. 2026). Similarly, in mice, paternal dietary perturbations alter the expression of sperm 5’ tDRs, and microinjection of these specific 5’ fragments into normal zygotes reprograms embryonic metabolic pathways and induces metabolic phenotypes in offspring (Chen et al. 2016). Although the molecular functions of cnidarian tDRs remain to be experimentally validated, the 5’-terminal dominance in sperm across three phylogenetically distinct cnidarians suggests that 5’-terminal biogenesis and paternal packaging of tDRs may be a deeply conserved metazoan trait that contributes to post-fertilization gene regulation.

Evaluation of specific isoacceptors revealed an accumulation of 5’ tDRs originating from the tRNA-GlyGCC isoacceptor across all three cnidarian species, a pattern that is mirrored in findings from bilaterians. In mammals, 5’ tDR-GlyGCC expression increases in response to a low-protein paternal diet and represses genes associated with the endogenous retroelement MERVL in early embryos (Sharma et al. 2016). Further, 5’ tDR-GlyGCC modulates *C. elegans* embryonic gene expression, including translation initiation factors, histones, and F-box genes, and inhibition of this specific fragment rescues offspring from heat sensitivity phenotypes (Galambos et al. 2026). The presence of 5’ tDR-GlyGCC in sperm across *A. hyacinthus, A. poculata,* and *N. vectensis* demonstrates that accumulation of specific and functional tDRs may be shared between early-diverging metazoans and bilaterians.

### Implications for paternal epigenetic inheritance

The identification of a diverse and non-random small RNA repertoire in three phylogenetically distinct cnidarian species (*Acropora hyacinthus, Astrangia poculata,* and *Nematostella vectensis*) provides the first empirical characterization of small RNAs in cnidarian gametes/sperm. The presence of functional small RNA classes (miRNAs, piRNAs, tDRs) demonstrates that cnidarian sperm carry a structured, non-random small RNA payload into fertilization. Further, the pathways that are putatively targeted by miRNAs and piRNAs point to conserved roles in cnidarians of regulating early post-fertilization gene expression. Establishing that early-diverging metazoans harbor these conserved small RNA classes and their potential functionality offers the first initial evidence for the molecular capacity of paternal epigenetic inheritance at the base of the animal tree.

We acknowledge that our findings are based on high-throughput sequencing and bioinformatic analyses; therefore, the functional roles of sperm small RNAs during fertilization and early development in cnidarians need to be experimentally validated. Further, whether these small RNA profiles actively respond to environmental shifts during gametogenesis and transmit these signals to offspring requires direct experimental manipulation. Nevertheless, the data presented here transforms theoretical discussions of cnidarian epigenetic inheritance into a testable framework, establishing a crucial baseline for future functional and evolutionary studies.

## Acknowledgements

We thank Hollie Putnam for *Acropora hyacinthus* sperm collections, as well as Alexandra Moen and Anya Hanson of the University of Rhode Island Diving and Research Program for support with *Astrangia poculata* collection. *A. hyacinthus* collections were completed under permits issued by the French Polynesian Government (Délégation à la Recherche) and the Hautcommissariat de la République en Polynésie Francaise (DTRT) (Protocole d’Accueil 2005-2023). *A. poculata* colonies were collected under Rhode Island Department of Environmental Management collector’s permit #970. The authors extend appreciation to the Temperate Coral Research Conferences hosted by Roger Williams University, Tufts University, and Southern Connecticut State University for fostering creative conversations leading to this work. The computational resources for this work were provided by the University of Rhode Island’s partnership with the Unity Research Computing Platform, a multi-institutional cluster led by the University of Massachusetts and the University of Rhode Island. This work was supported by NSF CAREER award OCE 2237658 to KLB and a Pew Biomedical Scholars Award to CCC. BHG was supported by NIH Predoctoral T32 HD083185 and NSF Ocean Sciences Postdoctoral Research Fellowship 2506815.

## Data availability

Data and code to reproduce the analyses presented here can be found at https://github.com/JillAshey/Cnidarian_sperm_smRNA and will be archived on Zenodo following manuscript acceptance. Sequence data is stored under NCBI BioProject PRJNA1491391.

## References

1. Admoni, Yael, Arie Fridrich, Paris K. Weavers, et al. 2025. “miRNA-Target Complementarity in Cnidarians Resembles Its Counterpart in Plants.” EMBO Reports 26 (3): 836–59. 10.1038/s44319-024-00350-z.

2. Akiyama, Yasutoshi, and Pavel Ivanov. 2023. “TRNA –derived RNAS: Biogenesis and Roles in Translational Control.” WIREs RNA 14 (6): e1805. 10.1002/wrna.1805.

3. Alexa, Adrian, and Jörg Rahnenführer. 2024. topGO: Enrichment Analysis for Gene Ontology. R. V. 2.62.0. Released. https://bioconductor.org/packages/topGO.

4. Altschul, Stephen F., Warren Gish, Webb Miller, Eugene W. Myers, and David J. Lipman. 1990. “Basic Local Alignment Search Tool.” Journal of Molecular Biology 215: 403–10. 10.1016/S0022-2836(05)80360-2.

5. Andrews, Simon R. 2010. FastQC: A Quality Control Tool for High Throughput Sequence Data. V. 0.12.1. Released. http://www.bioinformatics.babraham.ac.uk/projects/fastqc.

6. Ashey, Jill, and Katie L. Barott. 2026. “References Used for Cnidarian Sperm Small RNA Analysis.” Zenodo. 10.5281/zenodo.22772180.

7. Ashey, Jill, Chloé Gilligan, and Hollie M. Putnam. 2026. “A miRNA-Mediated Gene Regulatory Network Supports Seasonal Plasticity in the Temperate Coral Astrangia Poculata.” Journal of Experimental Biology, ahead of print. 10.1242/jeb.253001.

8. Ashey, Jill, Hollie M. Putnam, and M. Conor McManus. 2025. “Guided by the Northern Star Coral: A Research Synthesis and Roadmap for *Astrangia Poculata*.” Biology Letters 21 (3): 20240469. 10.1098/rsbl.2024.0469.

9. Ashey, Jill, Javier A. Rodriguez-Casariego, Kathleen M. Durkin, et al. 2026. “Characterization of Long Noncoding RNAs, microRNAs, and piwiRNAs in Reef-Building Corals.” Environmental Epigenetics 12 (1): dvag021. 10.1093/eep/dvag021.

10. Axtell, Michael J. 2013. “ShortStack: Comprehensive Annotation and Quantification of Small RNA Genes.” RNA 19 (6): 740–51. 10.1261/rna.035279.112.

11. Axtell, Michael J., Jakub O. Westholm, and Eric C. Lai. 2011. “Vive La Différence: Biogenesis and Evolution of microRNAs in Plants and Animals.” Genome Biology 12 (4): 221. 10.1186/gb-2011-12-4-221.

12. Barckmann, Bridlin, Stéphanie Pierson, Jérémy Dufourt, et al. 2015. “Aubergine iCLIP Reveals piRNA-Dependent Decay of mRNAs Involved in Germ Cell Development in the Early Embryo.” Cell Reports 12 (7): 1205–16. 10.1016/j.celrep.2015.07.030.

13. Baumgarten, Sebastian, Maha J. Cziesielski, Ludivine Thomas, et al. 2018. “Evidence for miNA-mediated Modulation of the Host Transcriptome in Cnidarian–Dinoflagellate Symbiosis.” Molecular Ecology 27 (2): 403–18. 10.1111/mec.14452.

14. Borbee, E. M., I. Changsut, J. S. McAlister, et al. 2026. “Development of Methods for Spawning, Fertilization, Larval Husbandry, and Settlement in the Temperate Coral Astrangia Poculata.” Frontiers in Marine Science 13 (August): 1901305. 10.3389/fmars.2026.1901305.

15. Chan, Patricia P., Brian Y. Lin, Allysia J. Mak, and Todd M. Lowe. 2021. “tRNAscan-SE 2.0: Improved Detection and Functional Classification of Transfer RNA Genes.” Nucleic Acids Research 49 (16): 9077–96. 10.1093/nar/gkab688.

16. Chang, Hao-Ming, Natalia J. Martinez, James E. Thornton, John P. Hagan, Khang D. Nguyen, and Richard I. Gregory. 2012. “Trim71 Cooperates with microRNAs to Repress Cdkn1a Expression and Promote Embryonic Stem Cell Proliferation.” Nature Communications 3 (1): 923. 10.1038/ncomms1909.

17. Chen, Peiwei, Alexei A. Kotov, Baira K. Godneeva, Sergei S. Bazylev, Ludmila V. Olenina, and Alexei A. Aravin. 2021. “piRNA-Mediated Gene Regulation and Adaptation to Sex-Specific Transposon Expression in *D. Melanogaster* Male Germline.” Genes & Development 35 (11–12): 914–35. 10.1101/gad.345041.120.

18. Chen, Qi, Menghong Yan, Zhonghong Cao, et al. 2016. “Sperm tsRNAs Contribute to Intergenerational Inheritance of an Acquired Metabolic Disorder.” Science 351 (6271): 397–400. 10.1126/science.aad7977.

19. Chen, Shifu, Yanqing Zhou, Yaru Chen, and Jia Gu. 2018. “Fastp: An Ultra-Fast All-in-One FASTQ Preprocessor.” Bioinformatics 34 (17): i884–90. 10.1093/bioinformatics/bty560.

20. Conine, Colin C., Fengyun Sun, Lina Song, Jaime A. Rivera-Pérez, and Oliver J. Rando. 2018. “Small RNAs Gained during Epididymal Transit of Sperm Are Essential for Embryonic Development in Mice.” Developmental Cell 46 (4): 470–480.e3. 10.1016/j.devcel.2018.06.024.

21. Conn, Trinity, Jill Ashey, Ross Cunning, and Hollie Putnam. 2025. “Genome Assembly and Annotation of *Acropora Pulchra* from Mo’orea, French Polynesia.” *Gigabyte*, ahead of print, March 28. 10.46471/gigabyte.153.

22. Crews, David, Ross Gillette, Samuel V. Scarpino, Mohan Manikkam, Marina I. Savenkova, and Michael K. Skinner. 2012. “Epigenetic Transgenerational Inheritance of Altered Stress Responses.” Proceedings of the National Academy of Sciences 109 (23): 9143–48. 10.1073/pnas.1118514109.

23. Czech, Benjamin, Marzia Munafò, Filippo Ciabrelli, et al. 2018. “piRNA-Guided Genome Defense: From Biogenesis to Silencing.” Annual Review of Genetics 52 (1): 131–57. 10.1146/annurev-genet-120417-031441.

24. Despard, Brecia A., Jason D. Selwyn, Allison N. Shupp, and Steven V. Vollmer. 2025. “A Network Approach to White Band Disease Challenged Staghorn Coral Acropora Cervicornis microRNAs and Their Targets.” Ecology and Evolution 15 (4): e71351. 10.1002/ece3.71351.

25. Ding, Jiandong, Shuigeng Zhou, and Jihong Guan. 2012. “Finding MicroRNA Targets in Plants: Current Status and Perspectives.” *Genomics*, Proteomics & Bioinformatics 10 (5): 264–75. 10.1016/j.gpb.2012.09.003.

26. Durkin, Kathleen M., Jill Ashey, Javier A. Rodriguez-Casariego, et al. 2026. “Cross-Talk among miRNAs, lncRNAs, and DNA Methylation in Three Coral Species Reveal Conserved Epigenetic Regulatory Architecture.” Preprint, bioRxiv. 10.64898/2026.07.19.739451.

27. Eklund, Anders, Paul Dufort, Mattias Villani, and Stephen LaConte. 2014. “BROCCOLI: Software for Fast fMRI Analysis on Many-Core CPUs and GPUs.” Frontiers in Neuroinformatics 8 (March). 10.3389/fninf.2014.00024.

28. Enright, Anton J., Bino John, Ulrike Gaul, Thomas Tuschl, Chris Sander, and Debora S. Marks. 2003. “MicroRNA Targets in Drosophila.” Genome Biology 5 (1): R1. 10.1186/gb-2003-5-1-r1.

29. Ewels, Philip, Måns Magnusson, Sverker Lundin, and Max Käller. 2016. “MultiQC: Summarize Analysis Results for Multiple Tools and Samples in a Single Report.” Bioinformatics 32 (19): 3047–48. 10.1093/bioinformatics/btw354.

30. Fiesinger, Anna, Abdoallah Sharaf, Rachel Alderdice, et al. 2026. “The Genome of the Reef-Building Coral Porites Harrisoni from the Southern Persian/Arabian Gulf.” Gigabyte 2026 (March): gigabyte174. 10.46471/gigabyte.174.

31. Fitz-James, Maximilian H., and Giacomo Cavalli. 2022. “Molecular Mechanisms of Transgenerational Epigenetic Inheritance.” Nature Reviews Genetics 23 (6): 325–41. 10.1038/s41576-021-00438-5.

32. Flynn, Jullien M., Robert Hubley, Clément Goubert, et al. 2020. “RepeatModeler2 for Automated Genomic Discovery of Transposable Element Families.” Proceedings of the National Academy of Sciences 117 (17): 9451–57. 10.1073/pnas.1921046117.

33. Gajigan, Andrian P., and Cecilia Conaco. 2017. “A microRNA Regulates the Response of Corals to Thermal Stress.” Molecular Ecology 26 (13): 3472–83. 10.1111/mec.14130.

34. Galambos, Nicholas S., Olivia J. Crocker, Blair K. Schneider, et al. 2026. “tRNA-Derived RNA Processing in Sperm Transmits Non-Genetically Inherited Phenotypes to Offspring in C. Elegans.” Nature Communications 17 (1): 3999. 10.1038/s41467-026-70029-7.

35. Gebert, Daniel, René F. Ketting, Hans Zischler, and David Rosenkranz. 2015. “piRNAs from Pig Testis Provide Evidence for a Conserved Role of the Piwi Pathway in Post-Transcriptional Gene Regulation in Mammals.” PLOS ONE 10 (5): e0124860. 10.1371/journal.pone.0124860.

36. Glass, Benjamin H., Jill Ashey, Amarachukwu R. Okongwu, Hollie M. Putnam, and Katie L. Barott. 2023. “Characterization of a Sperm Motility Signalling Pathway in a Gonochoric Coral Suggests Conservation across Cnidarian Sexual Systems.” Proceedings of the Royal Society B: Biological Sciences 290.

37. Glass, Benjamin H., Angela C. Ye, Cassidy N. Hemphill, Katelyn G. Jones, Anna G. Dworetzky, and Katie L. Barott. 2025. “Hypoxia Disrupts Sex-Specific Physiology and Gene Expression Leading to Decreased Fitness in the Estuarine Sea Anemone *Nematostella Vectensis*.” Molecular Ecology 34 (9): e17755. 10.1111/mec.17755.

38. Hausser, Jean, Afzal Pasha Syed, Biter Bilen, and Mihaela Zavolan. 2013. “Analysis of CDS-Located miRNA Target Sites Suggests That They Can Effectively Inhibit Translation.” Genome Research 23 (4): 604–15. 10.1101/gr.139758.112.

39. Ho, D. H., and W. W. Burggren. 2010. “Epigenetics and Transgenerational Transfer: A Physiological Perspective.” Journal of Experimental Biology 213 (1): 3–16. 10.1242/jeb.019752.

40. Hong, Zhidan, Sihan Huang, Li Li, et al. 2025. “Cracking the Code: How piRNA Pathway Shapes Spermatogenesis and Combats Male Infertility.” Frontiers in Cell and Developmental Biology 13 (September): 1657744. 10.3389/fcell.2025.1657744.

41. Hughes, Terry P., Michele L. Barnes, David R. Bellwood, et al. 2017. “Coral Reefs in the Anthropocene.” Nature 546 (7656): 82–90. 10.1038/nature22901.

42. Iwasaki, Yuka W., Kensaku Murano, Hirotsugu Ishizu, et al. 2016. “Piwi Modulates Chromatin Accessibility by Regulating Multiple Factors Including Histone H1 to Repress Transposons.” Molecular Cell 63 (3): 408–19. 10.1016/j.molcel.2016.06.008.

43. Jehn, Julia, Daniele Gerbert, Frank Pipilescu, et al. 2018. “PIWI Genes and piRNAs Are Ubiquitously Expressed in Mollusks and Show Patterns of Lineage-Specific Adaptation.” Communications Biology 1.

44. Juliano, Celina E., Adrian Reich, Na Liu, et al. 2014. “PIWI Proteins and PIWI-Interacting RNAs Function in *Hydra* Somatic Stem Cells.” Proceedings of the National Academy of Sciences 111 (1): 337–42. 10.1073/pnas.1320965111.

45. Kaikkonen, M. U., M. T. Y. Lam, and C. K. Glass. 2011. “Non-Coding RNAs as Regulators of Gene Expression and Epigenetics.” Cardiovascular Research 90 (3): 430–40. 10.1093/cvr/cvr097.

46. Kayal, Ehsan, Bastian Bentlage, M. Sabrina Pankey, et al. 2018. “Phylogenomics Provides a Robust Topology of the Major Cnidarian Lineages and Insights on the Origins of Key Organismal Traits.” BMC Evolutionary Biology 18 (1): 68. 10.1186/s12862-018-1142-0.

47. Kopylova, Evguenia, Laurent Noé, and Hélène Touzet. 2012. “SortMeRNA: Fast and Accurate Filtering of Ribosomal RNAs in Metatranscriptomic Data.” Bioinformatics 28 (24): 3211–17. 10.1093/bioinformatics/bts611.

48. Krawetz, Stephen A. 2005. “Paternal Contribution: New Insights and Future Challenges.” Nature Reviews Genetics 6 (8): 633–42. 10.1038/nrg1654.

49. Krishna, Srikar, Aparna Nair, Sirisha Cheedipudi, et al. 2013. “Deep Sequencing Reveals Unique Small RNA Repertoire That Is Regulated during Head Regeneration in Hydra Magnipapillata.” Nucleic Acids Research 41 (1): 599–616. 10.1093/nar/gks1020.

50. Langmead, Ben, and Steven L. Salzberg. 2012. “Fast Gapped-Read Alignment with Bowtie 2.” Nature Methods 9: 357–59. 10.1038/nmeth.1923.

51. Lee, Grace S., and Colin C. Conine. 2022. “The Transmission of Intergenerational Epigenetic Information by Sperm microRNAs.” Epigenomes 6 (2): 12. 10.3390/epigenomes6020012.

52. Lee, Grace S., James Garifallou, Samantha L. Higgins, et al. 2026. “Physiologic Variation in Sperm miRNAs Tune Embryonic Gene Regulatory Programs and Developmental Outcomes.” Preprint, bioRxiv. 10.64898/2026.03.04.709527.

53. López-Nandam, Elora H., Rebecca Albright, Erik A. Hanson, Elizabeth A. Sheets, and Stephen R. Palumbi. 2023. “Mutations in Coral Soma and Sperm Imply Lifelong Stem Cell Renewal and Cell Lineage Selection.” Proceedings of the Royal Society B: Biological Sciences 290 (20221766). 10.1098/rspb.2022.1766.

54. Mehta, Poonam, and Rajender Singh. 2025. “The Composition of Human Sperm sncRNAome: A Cross-Country Small RNA Profiling.” Reproductive Biology and Endocrinology 23 (1): 36. 10.1186/s12958-025-01358-3.

55. Miller, D., E. Ball, and U. Technau. 2005. “Cnidarians and Ancestral Genetic Complexity in the Animal Kingdom.” Trends in Genetics 21 (10): 536–39. 10.1016/j.tig.2005.08.002.

56. Moran, Yehu, David Fredman, Daniela Praher, et al. 2014. “Cnidarian microRNAs Frequently Regulate Targets by Cleavage.” Genome Research 24 (4): 651–63. 10.1101/gr.162503.113.

57. Mu, Jian, Zhou Zhou, Qing Sang, and Lei Wang. 2022. “The Physiological and Pathological Mechanisms of Early Embryonic Development.” Fundamental Research 2 (6): 859–72. 10.1016/j.fmre.2022.08.011.

58. Muthukumar, Sowndarya, Cai-Tao Li, Ru-Juan Liu, and Cristian Bellodi. 2024. “Roles and Regulation of tRNA-Derived Small RNAs in Animals.” Nature Reviews Molecular Cell Biology 25 (5): 359–78. 10.1038/s41580-023-00690-z.

59. Naveed, Muhammad, Zhaokang Shen, and Jianqiang Bao. 2025. “Sperm-Borne Small Non-Coding RNAs: Potential Functions and Mechanisms as Epigenetic Carriers.” Cell & Bioscience 15 (1): 5. 10.1186/s13578-025-01347-4.

60. Noel, Benjamin, France Denoeud, Alice Rouan, et al. 2023. “Pervasive Tandem Duplications and Convergent Evolution Shape Coral Genomes.” Genome Biology 24 (1): 123. 10.1186/s13059-023-02960-7.

61. Norouzitallab, Parisa, Kartik Baruah, Daisy Vanrompay, and Peter Bossier. 2019. “Can Epigenetics Translate Environmental Cues into Phenotypes?” Science of The Total Environment 647 (January): 1281–93. 10.1016/j.scitotenv.2018.08.063.

62. Obura, David O., Beth Polidoro, Francoise Cabada-Blanco, et al. 2026. “Nearly Half of Reef-Building Corals Face Elevated Relative Risk of Extinction.” *Nature*, ahead of print, October 15. 10.1038/s41586-026-11263-3.

63. Perillo, Giulia, Keigo Shibata, and Pei-Hsuan Wu. 2023. “piRNAs in Sperm Function and Embryo Viability.” Reproduction 165 (3): R91–102. 10.1530/REP-22-0312.

64. Pezic, Dubravka, Sergei A. Manakov, Ravi Sachidanandam, and Alexei A. Aravin. 2014. “piRNA Pathway Targets Active LINE1 Elements to Establish the Repressive H3K9me3 Mark in Germ Cells.” Genes & Development 28 (13): 1410–28. 10.1101/gad.240895.114.

65. Praher, Daniela, Bob Zimmermann, Rohit Dnyansagar, et al. 2021. “Conservation and Turnover of miRNAs and Their Highly Complementary Targets in Early Branching Animals.” Proceedings of the Royal Society B: Biological Sciences 288 (1945): 20203169. 10.1098/rspb.2020.3169.

66. Praher, Daniela, Bob Zimmermann, Grigory Genikhovich, et al. 2017. “Characterization of the piRNA Pathway during Development of the Sea Anemone *Nematostella Vectensis*.” RNA Biology 14 (12): 1727–41. 10.1080/15476286.2017.1349048.

67. Quinlan, Aaron R., and Ira M. Hall. 2010. “BEDTools: A Flexible Suite of Utilities for Comparing Genomic Features.” Bioinformatics 26 (6): 841–42. 10.1093/bioinformatics/btq033.

68. R Core Team. 2024. R: A Language and Environment for Statistical Computing. R Foundation for Statistical Computing, released. https://www.R-project.org/.

69. Rodgers, Ali B., Christopher P. Morgan, Stefanie L. Bronson, Sonia Revello, and Tracy L. Bale. 2013. “Paternal Stress Exposure Alters Sperm MicroRNA Content and Reprograms Offspring HPA Stress Axis Regulation.” The Journal of Neuroscience 33 (21): 9003–12. 10.1523/JNEUROSCI.0914-13.2013.

70. Rodgers, Ali B., Christopher P. Morgan, N. Adrian Leu, and Tracy L. Bale. 2015. “Transgenerational Epigenetic Programming via Sperm microRNA Recapitulates Effects of Paternal Stress.” Proceedings of the National Academy of Sciences 112 (44): 13699–704. 10.1073/pnas.1508347112.

71. Rojas-Ríos, Patricia, and Martine Simonelig. 2018. “piRNAs and PIWI Proteins: Regulators of Gene Expression in Development and Stem Cells.” Development 145 (17): dev161786. 10.1242/dev.161786.

72. Rosenkranz, David, Chung-Ting Han, Elke F. Roovers, Hans Zischler, and René F. Ketting. 2015. “Piwi Proteins and piRNAs in Mammalian Oocytes and Early Embryos: From Sample to Sequence.” Genomics Data 5 (September): 309–13. 10.1016/j.gdata.2015.06.026.

73. Rouget, Christel, Catherine Papin, Anthony Boureux, et al. 2010. “Maternal mRNA Deadenylation and Decay by the piRNA Pathway in the Early Drosophila Embryo.” Nature 467 (7319): 1128–32. 10.1038/nature09465.

74. Seemann, Torsten. 2013. Barrnap 0.9: Rapid Ribosomal RNA Prediction. V. 0.9. Released. http://github.com/tseemann/barrnap.

75. Shahid, Saima, and Michael J. Axtell. 2014. “Identification and Annotation of Small RNA Genes Using ShortStack.” Methods 67 (1): 20–27. 10.1016/j.ymeth.2013.10.004.

76. Sharma, Upasna. 2019. “Paternal Contributions to Offspring Health: Role of Sperm Small RNAs in Intergenerational Transmission of Epigenetic Information.” Frontiers in Cell and Developmental Biology 7 (October): 215. 10.3389/fcell.2019.00215.

77. Sharma, Upasna, Colin C. Conine, Jeremy M. Shea, et al. 2016. “Biogenesis and Function of tRNA Fragments during Sperm Maturation and Fertilization in Mammals.” Science 351 (6271): 391–96. 10.1126/science.aad6780.

78. Siomi, Mikiko C., Kaoru Sato, Dubravka Pezic, and Alexei A. Aravin. 2011. “PIWI-Interacting Small RNAs: The Vanguard of Genome Defence.” Nature Reviews Molecular Cell Biology 12 (4): 246–58. 10.1038/nrm3089.

79. Smit, A. F. A., Robert Hubley, and P. Green. 2015. RepeatMasker Open-4.0. V. 4.0. Released. https://www.repeatmasker.org/.

80. Stankiewicz, Kathryn H., Nadége Guiglielmoni, Sheila A. Kitchen, et al. 2025. “Genomic Comparison of the Temperate Coral Astrangia Poculata with Tropical Corals Yields Insights into Winter Quiescence, Innate Immunity, and Sexual Reproduction.” G3 Genes|Genomes|Genetics jkaf033. 10.1093/g3journal/jkaf033.

81. Su, Zhangli, Briana Wilson, Pankaj Kumar, and Anindya Dutta. 2020. “Noncanonical Roles of tRNAs: tRNA Fragments and Beyond.” Annual Review of Genetics 54 (1): 47–69. 10.1146/annurev-genet-022620-101840.

82. Tadros, Wael, and Howard D. Lipshitz. 2009. “The Maternal-to-Zygotic Transition: A Play in Two Acts.” Development 136 (18): 3033–42. 10.1242/dev.033183.

83. Teefy, Bryan B., Stefan Siebert, Jack F. Cazet, Haifan Lin, and Celina E. Juliano. 2020. “PIWI–piRNA Pathway-Mediated Transposable Element Repression in Hydra Somatic Stem Cells.” RNA 26: 550–63.

84. Tomar, A., M. Gomez-Velazquez, R. Gerlini, et al. 2024. “Epigenetic Inheritance of Diet-Induced and Sperm-Borne Mitochondrial RNAs.” Nature 630 (8017): 720–27. 10.1038/s41586-024-07472-3.

85. Torda, Gergely, Jennifer M. Donelson, Manuel Aranda, et al. 2017. “Rapid Adaptive Responses to Climate Change in Corals.” Nature Climate Change 7 (9): 627–36. 10.1038/nclimate3374.

86. Torres-Fernández, Lucia A., Jana Emich, Yasmine Port, et al. 2021. “TRIM71 Deficiency Causes Germ Cell Loss During Mouse Embryogenesis and Is Associated With Human Male Infertility.” Frontiers in Cell and Developmental Biology 9 (May): 658966. 10.3389/fcell.2021.658966.

87. Trigg, Natalie A., and Colin C. Conine. 2024. “Epididymal Acquired Sperm microRNAs Modify Post-Fertilization Embryonic Gene Expression.” Cell Reports 43 (9): 114698. 10.1016/j.celrep.2024.114698.

88. Trigg, Natalie, John E. Schjenken, Jacinta H. Martin, et al. 2024. “Subchronic Elevation in Ambient Temperature Drives Alterations to the Sperm Epigenome and Accelerates Early Embryonic Development in Mice.” Proceedings of the National Academy of Sciences 121 (47): e2409790121. 10.1073/pnas.2409790121.

89. Vastenhouw, Nadine L., Wen Xi Cao, and Howard D. Lipshitz. 2019. “The Maternal-to-Zygotic Transition Revisited.” Development 146 (11): dev161471. 10.1242/dev.161471.

90. Wu, Pei-Hsuan, Yu Fu, Katharine Cecchini, et al. 2020. “The Evolutionarily Conserved piRNA-Producing Locus Pi6 Is Required for Male Mouse Fertility.” Nature Genetics 52 (7): 728–39. 10.1038/s41588-020-0657-7.

91. Zimmermann, Bob, Juan D. Montenegro, Sofia M. C. Robb, et al. 2023. “Topological Structures and Syntenic Conservation in Sea Anemone Genomes.” Nature Communications 14 (1): 8270. 10.1038/s41467-023-44080-7.

